# Stearoyl-CoA desaturase mediated monounsaturated fatty acid availability supports humoral immunity

**DOI:** 10.1101/2020.04.22.028613

**Authors:** Xian Zhou, Xingxing Zhu, Chaofan Li, Yanfeng Li, Zhenqing Ye, Virginia Shapiro, John A. Copland, Taro Hitosugi, David Bernlohr, Jie Sun, Hu Zeng

## Abstract

Immune cells can metabolize glucose, amino acids, and fatty acids (FAs) to generate energy. The role of different FA species, and their impacts on humoral immunity remains poorly understood. Here we report that proliferating B cells require monounsaturated FAs (MUFA) to maintain mitochondrial metabolism and mTOR activity, and to prevent excessive autophagy and endoplasmic reticulum (ER) stress. Furthermore, B cell extrinsic Stearoyl-CoA desaturase (SCD) activity generates MUFA to support early B cell development and germinal center (GC) formation *in vivo* during immunization and influenza infection. Thus, SCD-mediated MUFA production is critical for humoral immunity.

## INTRODUCTION

There is growing evidence that B lymphocyte development and activation are regulated by metabolic processes^1^. In particular, glucose metabolism was shown to be activated by BCR or TLR ligand stimulation and support B cell function^2–4^. Recent studies also demonstrated that activation of mitochondrial oxidative phosphorylation by TLR ligand or CD40L, and supported by glutamine import, is required for B cell survival^5,6^. Consistent with these findings, the central metabolic regulator, mammalian target of rapamycin (mTOR), is key to support B cell development and humoral response, through diverse metabolic actions, including organelle biogenesis and various anabolic processes^7–10^. However, a recent study suggested that activated B cells utilize glucose mainly for ribonucleotide and FA biosynthesis, but not for lactate production or feeding into TCA cycle^6^. Thus, biomass accumulation, including FA biosynthesis, appears to be the main features of early B cell activation^11^. Glucose, glutamine and FAs are the three major carbon sources for most cell types. While regulation of glucose and glutamine metabolism has been extensively studied in immunometabolism field, the contribution of different FA species to B cell function remains poorly understood.

It is long recognized that both malnutrition and obesity impair humoral immunity^12,13^. Because a plethora of nutrients and metabolites are altered in general malnutrition or obesity, it is challenging to parse how each metabolite, especially different fatty acids, impacts humoral immunity and thus exact mechanisms linking immunity to systemic metabolism remains obscure. A recent study showed that germinal center (GC) B cells mainly utilize FA oxidation, rather than glycolysis, to meet their energetic need, highlighting the importance of FA availability for humoral immunity^14^. While FAs are known to contribute to energy metabolism through β-oxidation, most studies in immunometabolism do not distinguish different FAs. Therefore, it is unclear how different FA species may contribute to immunity. To date, most studies on lipid metabolism and immune function have focused on the relationship between different types of diets with varying FA contents and systemic inflammation^15^, whereas how endogenously generated FA species impact humoral immunity remains unknown.

Stearoyl-CoA desaturase (SCD) is a rate limiting enzyme in *de novo* FA biosynthesis. It converts saturated FA (SFA) into mono-unsaturated FAs (MUFAs), including oleic acid (OA) and palmitoleic acid (PO). SCD plays a central role in fuel metabolism and constitutes a potential therapeutic target for treatment of obesity and cancer ^16^. SCD1 deficient mice are protected from high-fat-diet and high-carbohydrate-diet induced obesity and hepatic steatosis^17,18^. Interestingly, despite ready access to dietary sources of OA, some of the metabolic defects in SCD1 deficient mice persist, even when the mice were fed with diet containing high level of OA^18–20^, highlighting the importance of endogenously synthesized MUFA for proper cellular function and FA metabolism.

Here, we present mechanistic evidence that B cell development and activation require SCD generated MUFA, particularly OA, which maintains B cell metabolic fitness partly by supporting mitochondrial oxidative phosphorylation and mTORC1 activity, and preventing excessive autophagy and ER stress. *In vivo*, B cells mainly rely on B cell-extrinsic SCD activity to provide MUFAs. In response to immune challenges, the host enhances MUFA availability, partly through SCD activity, which is required to sustain antibody production. Suppression of SCD reduces humoral immune response to immunization, and weakens immune defense against respiratory influenza A virus infection. Thus, our results provide a novel link between endogenous biosynthesis of a specific FA specie to humoral immunity in immunization and anti-influenza immune defense.

## RESULTS

### SCD mediated MUFA biosynthesis is activated during B cell activation

We first sought to determine the FA biosynthesis gene expression program during B cell activation. RNA sequencing was performed using fresh murine B cells and LPS/IL-4 activated B cells. Major genes involved in FA biosynthesis, including *Acaca*, *Elovl1, Elovl5, Elovl6, Fasn* and *Scd2*, were all increased (Fig. 1A). Among the 4 murine *Scd* genes, *Scd3* and *Scd4* were below detection limit. *Scd1* expression was slightly reduced, while *Scd2* expression was substantially increased (Fig. 1B). The increase of *Scd2* expression was also confirmed by immunoblot (Fig. 1C). A similar dynamic occurred during the first 24 hours of activation, in which *Scd2* expression showed the most upregulation after 24 hours stimulation (Fig. S1A). The induction of SCD2 was not specific to LPS/IL-4 stimulation, because anti-IgM, anti-CD40 and CpG, but not IL-4 alone, also induced SCD2 protein expression at 24 hours after activation (Fig. S1B). Furthermore, activation of human B cells also induced robust *SCD* expression (Fig. 1D). Finally, the upregulation of FA biosynthesis genes was dependent on mTORC1 signaling, as rapamycin treatment blocked their increased expression (Fig. 1E), consistent with the anabolic function of mTORC1 on lipid biosynthesis. Thus, antigenic stimulation of B cells activates FA biosynthetic pathway in an mTORC1 dependent manner.

**Figure 1.**
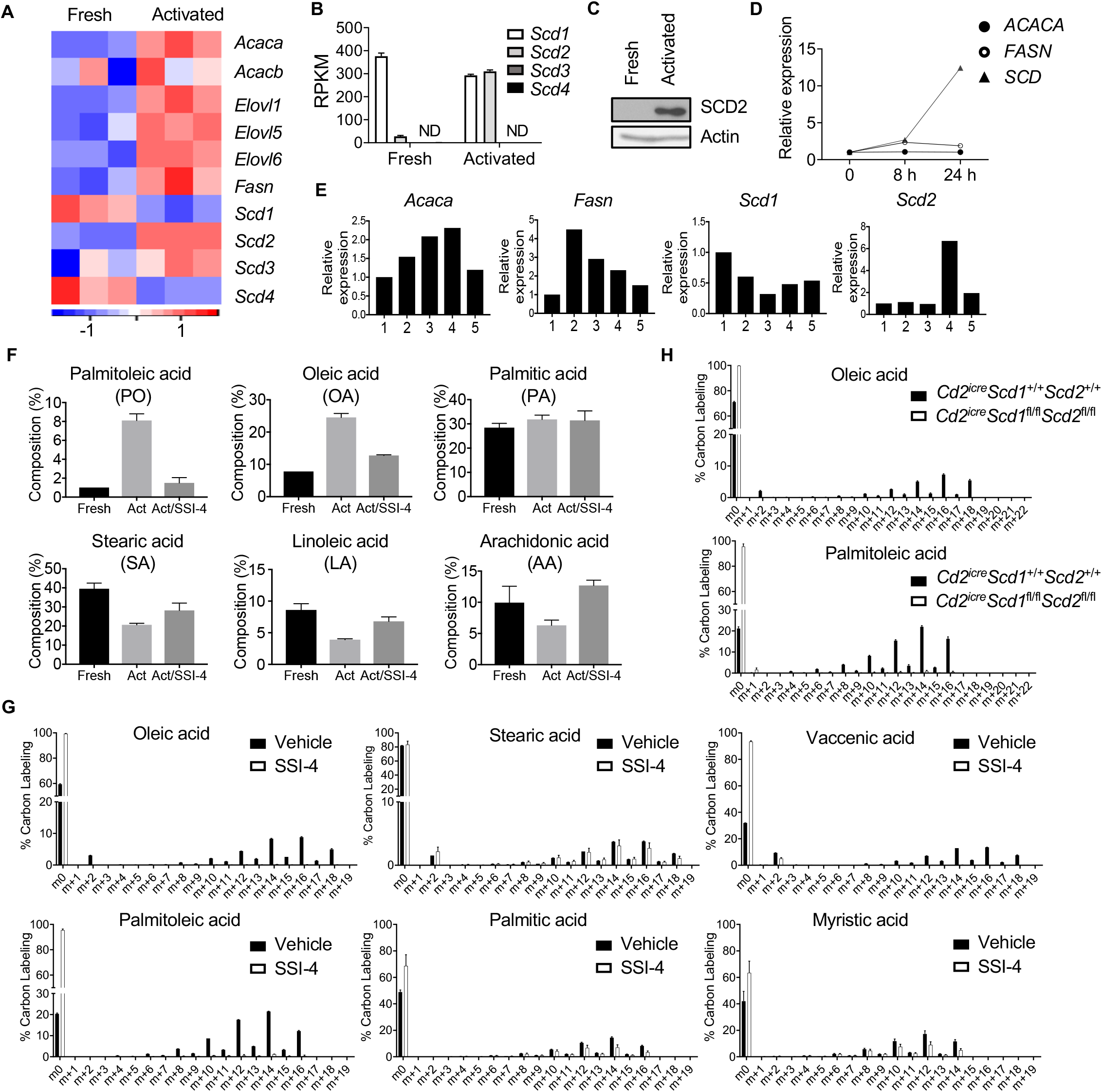
SCD mediated MUFA biosynthesis during B cell activation *in vitro.* (A) Heat map of genes expression related to fatty acid synthesis in murine fresh and activated B cells with LPS/IL-4. (B) mRNA expression of 4 *Scd* isoforms’ expression in freshly isolated or activated B cells, extracted from RNAseq data. “ND”, Not detected. (C) Immunoblot analysis of SCD2 in murine fresh and activated B cells by immunoblot. (D) Reverse transcription polymerase chain reaction (RT-PCR) analysis of *ACACA*, *FASN,* and *SCD* expression in human B cells at 0, 8 and 24 h during activation with CpG/anti-CD40/IL-15/IL-10/IL-2. (E) RT-PCR analysis of *Acaca*, *Fasn*, *Scd1*, and *Scd2* expression in murine B cells with or without rapamycin (10 nM) treatment at 8 and 24 h. 1, fresh B cell; 2, B cells activated with LPS/IL-4 for 8 h in vitro, 3, B cells activated with LPS/IL-4 plus rapamycin for 8 h; 4, B cells activated with LPS/IL-4 for 24 h; 5, B cell activated with LPS/IL-4 plus rapamycin treatment for 24 h. (F) Composition of palmitoleic acid (PO), oleic acid (OA), palmitic acid (PA), stearic acid (SA), linoleic acid (LA), and arachidonic acid (AA) in fresh and activated B cells treated with vehicle or 100 nM SCD inhibitor SSI-4 for 48 h was measured by LC-MS/MS. (G) Isotopomer distribution for ^13^C labeled glucose in fatty acid synthesis metabolites OA, SA, vaccenic acid (VA), PO, PA and myristic acid (MA) in activated B cell treated with vehicle or SSI-4 inhibitor at 48 h. (H) Comparison of ^13^C labeled glucose incorporation in OA and PO in activated B cells isolated from either WT or *Cd2^iCre^Scd1*^fl/fl^*Scd2*^fl/fl^ mice.

To further examine the functional outcome of increased FA biosynthesis gene expression, we employed triple quadrupole liquid chromatography/tandem mass spectrometry (LC-MS/MS) to examine specific FA content in fresh and activated B cells. Our targeted metabolomics showed that PO and OA, the main MUFAs produced by SCD, exhibited the highest increase of their relative contents in activated B cells compared to unstimulated B cells. The composition of palmitic acid (PA), the precursor of PO, did not change substantially, while the composition of stearic acid (SA), the precursor of OA and the PUFAs, including linoleic acid (LA) and arachidonic acid (AA), modestly reduced after activation (Fig. 1F). This dramatic accumulation of MUFAs was dependent on SCD activity, because treatment with an SCD specific inhibitor, SSI-4^21^, largely reversed it (Fig. 1F). Furthermore, the *de novo* biosynthesis of FAs from glucose was confirmed in a U-^13^C-6-glucose tracing experiment. When B cells were activated in medium containing uniformly ^13^C labeled glucose, our LC-MS/MS assay detected substantial ^13^C incorporation in OA, PO and vaccenic acid (VA, another MUFA), which were all abolished when cells were treated with SSI-4. However, the incorporation of ^13^C into SA, PA and myristic acid (MA) was largely unaffected by SSI-4 treatment (Fig. 1G). These observations were further confirmed using a genetic model, in which *Scd1* and *Scd2* were deleted in B cells through *Cd2*iCre, an optimized variant of Cre recombinase under human *CD2* promoter and locus control region that leads to efficient recombination in lymphocytes^22,23^. SCD1 and SCD2 deficiency completely eliminated ^13^C-glucose incorporation into OA, PO and VA, but did not substantially affect ^13^C incorporation into PA, SA, or MA (Fig. 1H, and Fig. S1C). Incorporation of ^13^C into other PUFAs, including arachidonic acid, linoleic acid and α-linolenic acid, was not detected in our assays (data not shown), suggesting that B cells do not have the capacity to generate these FAs from glucose *de novo*. Therefore, these results showed that B cell activation is associated with activation of SCD activity and increased proportion of SCD-generated MUFA content.

### SCD-generated MUFA supports B cell proliferation and class switch in vitro

To further investigate the impact of SCD generated MUFA on B cell functions, we measured the proliferation of mouse B cells activated with LPS/IL-4 in the presence of different SCD inhibitors, including SSI-4, MF438, and A939572. All were capable of inhibiting B cell proliferation (Fig. 2A). Importantly, exogenous OA was able to rescue the proliferation defects caused by SCD inhibition, and promote cell number accumulation, demonstrating that the enzymatic activity of SCD is required for B cell proliferation (Fig. 2B, C). Similar phenotypes were observed when we stimulated B cells with anti-IgM/anti-CD40 or TLR9 ligand, CpG (Fig. 2B). LPS/IL-4 also stimulates class switch to IgG1. We found that SCD inhibition strongly suppressed IgG1 class switch (Fig. 2D). Exogenous OA alone further enhanced B cell proliferation (measured by cell number (Fig. 2C)) and IgG1 class switch, and it could fully restore both parameters upon SSI-4 treatment (Fig. 2C, D). In contrast, PO alone did not improve class switch and had a substantial, but incomplete, rescue effects on proliferation and IgG1 class switch upon SSI-4 treatment. PA and SA showed no rescue effects upon SSI-4 treatment (Fig. 2D). Moreover, unlike OA, supply of exogenous PA and SA were unable to promote the proliferation and class switch (Fig. S2A). Of note, the concentration of exogenous FAs we applied in these *in vitro* assays was based on the levels of serum non-esterified fatty acids (NEFAs) in WT mice (Table S1). These results suggest that proliferating B cells preferentially utilize MUFA, especially OA, rather than SFAs. Moreover, we isolated splenic B cells from *Scd1* and *Scd2* deficient mice and stimulated them with LPS/IL-4. SCD1/2 deficient B cells proliferated poorly, had highly reduced IgG1 class switch compared to those from control mice. Exogenous OA fully restored proliferation and IgG1 class switch, while PO had a partial rescue effect (Fig. 2E). Lastly, we confirmed these observations using human B cells isolated from peripheral blood mononuclear cells (PBMC) of healthy donors. Exogenous OA promoted human B cell proliferation (Fig. 2F) and activation measured by expression of activation marker CD27 (Fig. 2G), and it also restored B cell proliferation and CD27 expression upon SSI-4 treatment (Fig. 2F and 2G). Thus, MUFAs, particularly OA, promote B cell proliferation and class switch *in vitro*.

**Figure 2.**
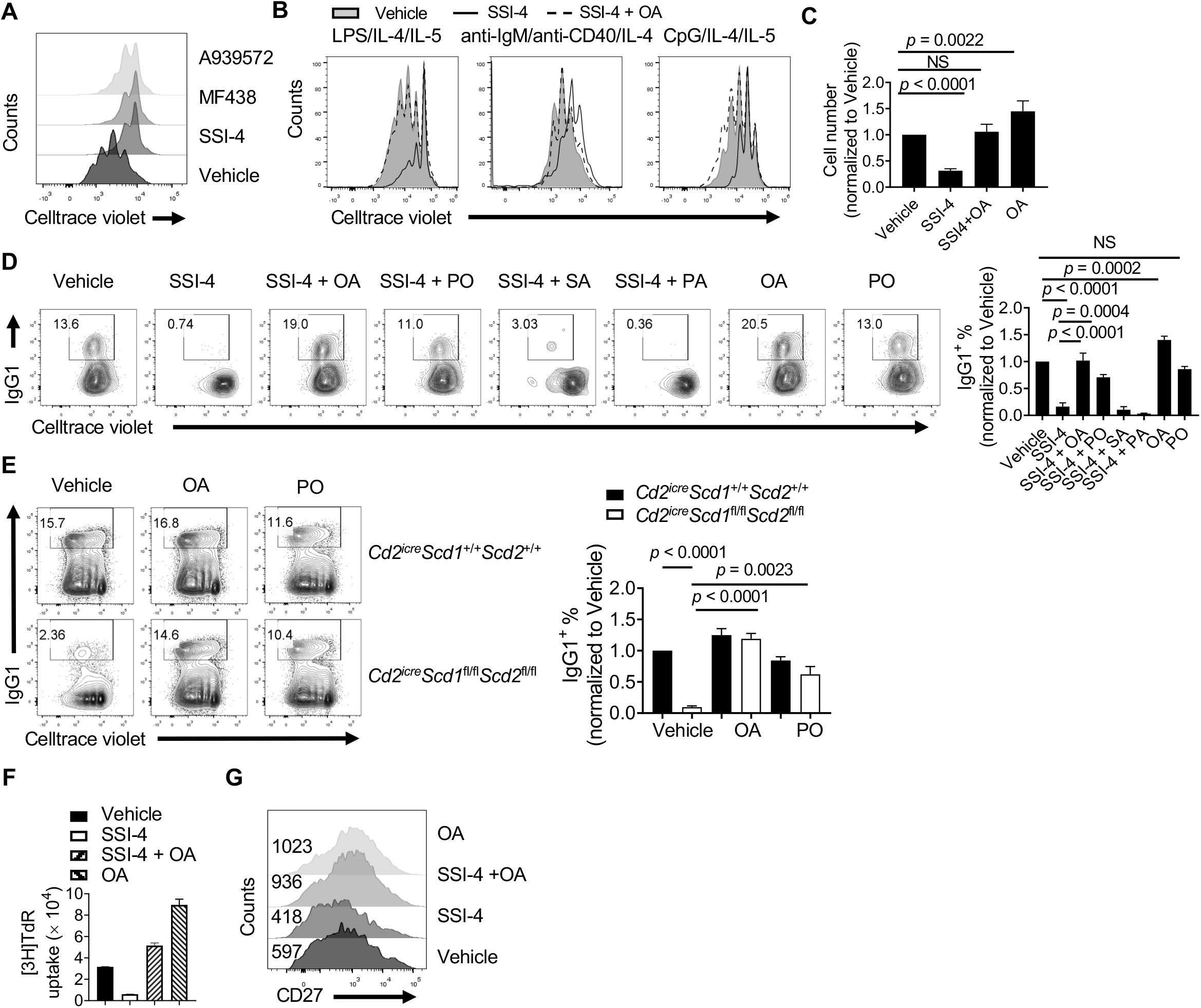
SCD-generated MUFA is required for B cell proliferation and class switch *in vitro*. (A) Cell proliferation measured by dilution of Celltrace Violet (CTV) dye in LPS/IL-4 activated murine B cell treated with SCD inhibitors A939572 (1 μM), MF438 (1 μM), and SSI-4 (1 μM), respectively. (B) Cell proliferation of murine B cells stimulated with LPS/IL-4/IL-5, anti-IgM/anti-CD40/IL-4, or CpG/IL-4/IL-5 in presence of vehicle, 100 nM SSI-4, or SSI-4 plus exogenous 100 μM OA, respectively for 3 days. (C) Murine B cells were stimulated with LPS/IL-4/IL-5 for 3 days in the presence of indicated inhibitor and/or fatty acids. Numbers of activated B cells were summarized, and were normalized to vehicle group. (D) Flow cytometry analysis of murine B cell proliferation and class switch to IgG1 activated with LPS/IL-4 in presence of vehicle, SCD inhibitor SSI-4, SSI-4 with exogenous OA (100 μM), SSI-4 with exogenous PO (25 μM), SSI-4 with exogenous SA (25 μM), SSI-4 with exogenous PA (100 μM), OA alone, and PO alone. Right, summary of IgG1^+^ B cell percentages normalized against vehicle treated samples. (E) Cell proliferation and class switch to IgG1 in activated B cells isolated from WT or *Cd2^iCre^Scd1*^fl/fl^*Scd2*^fl/fl^ mice detected by flow cytometry. Right, summary of IgG1^+^ B cell percentage normalized to vehicle treated WT samples. (F) Human naive B cell proliferation was measured by thymidine incorporation after activation in the presence of vehicle, SCD inhibitor SSI-4 (10 nM), SSI-4 plus exogenous OA, and OA alone. (G) Flow cytometry analysis of CD27 expression on activated human B cells in the presence of vehicle, SSI-4, SSI-4 plus OA or OA alone. Numbers indicate the mean fluorescence intensity (MFI). *p* values were calculated with one-way ANOVA. NS, not significant. Data were representative of at least 3 (A-E) or 2 (F and G) independent experiments. Error bars represent SEM.

### MUFA maintains B cell metabolic fitness

B cell activation is accompanied by increased oxygen consumption, which are important to support humoral immunity^2,5,6^. FAs can be utilized to fuel mitochondrial oxidative phosphorylation (OXPHOS) and to provide energy, but it is unclear how different FAs contribute to B cell metabolism. Indeed, the inhibitor etomoxir, which blocks carnitine palmitoyltransferase I (CPT1)-mediated FA import to mitochondrial, reduced B cell class switch, and negated the effects of OA treatment at 40 μM, a dose selective to CPT1 inhibition^14^ (Fig. 3A). This data indicated that fatty acid oxidation (FAO) of OA supports B cell function. To analyze the metabolic function of SFAs and MUFAs on B cell OXPHOS, we measured the oxygen consumption rate (OCR) on B cells stimulated by CpG/IL-4/IL-5 in the presence of different exogenous FAs. Addition of OA, but not SFA, enhanced B cell respiration (Fig. 3B). Similar findings were observed using LPS/IL-4 and anti-IgM/IL-4 stimulation (Fig. S3A, S3B). The augmented respiration was also observed in activated human B cells supplied with OA (Fig. 3C). OA mediated increase of respiration was dose dependent (Fig. 3D). The effect of PO on B cell metabolism appeared to be highly variable. While it did not promote respiration upon CpG and anti-IgM/anti-CD40 stimulation (Fig. 3B, Fig. S3B), it did enhance OCR upon LPS/IL-4 stimulation in mouse B cells (Fig. S3A) and in human B cells (Fig. 3C). Moreover, OA and, to a less degree, PO, increased the glycolytic capacity of activated murine B cells while SFA had either negative or no effects on glycolysis (Fig. 3E). Consistent with murine data, OA improved human B cell glycolysis (Fig. 3F). Thus, our data indicate that OA promotes both mouse and human B cell OXPHOS and glucose metabolism. Conversely, inhibition of SCD activity via SSI-4 strongly suppressed B cell respiration and glycolysis (Fig. 3G). Consistent with defective energetic and anabolic metabolism, we observed reduced mitochondrial membrane potential measured by staining of tetramethylrhodamine (TMRM) when SCD activity was inhibited, which was corrected with exogenous OA (Fig. 3H). OA alone was also able to increase mitochondrial membrane potential (Fig. 3H). These data suggest that provision of OA through SCD activity is critical for B cell metabolic fitness.

**Figure 3.**
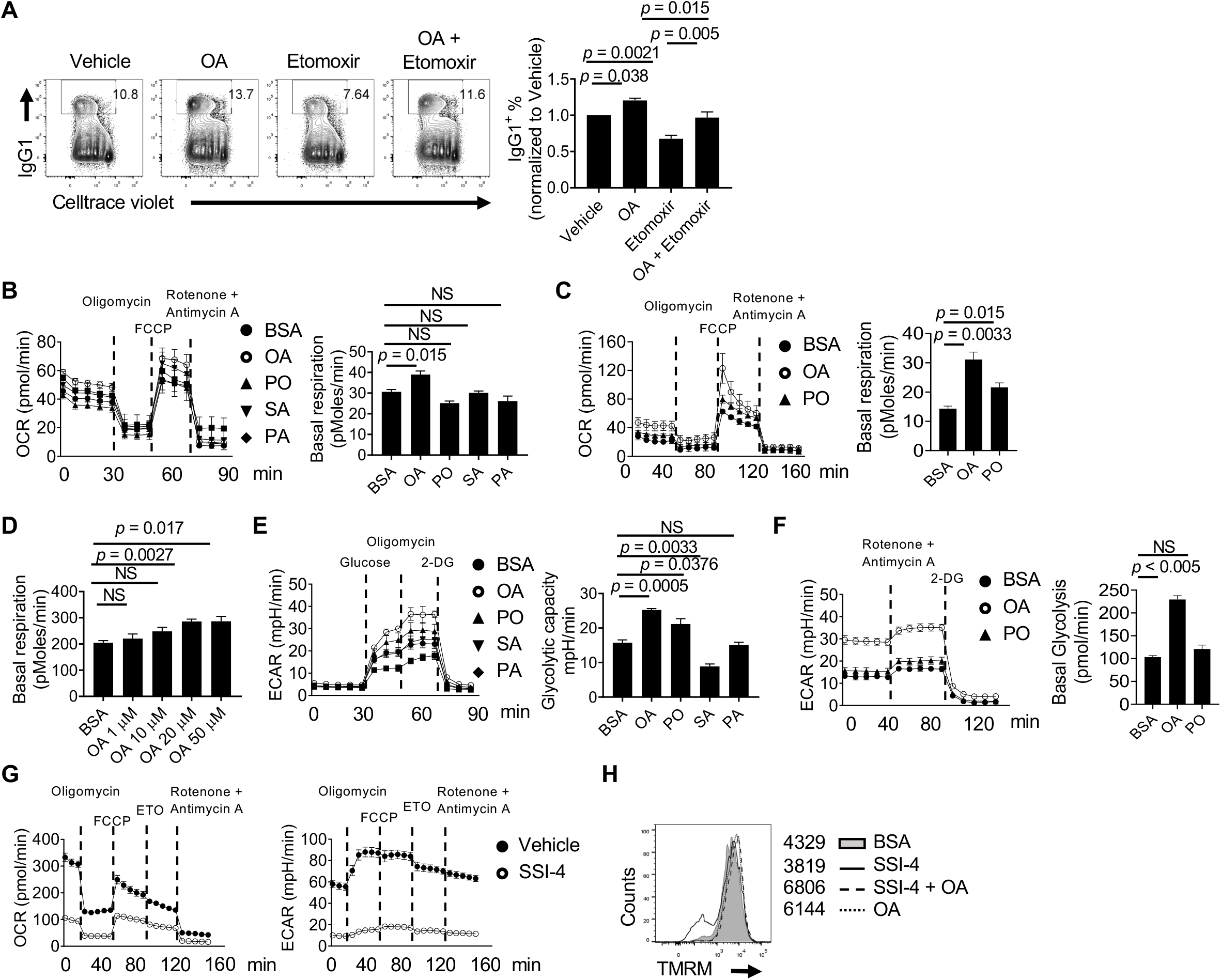
MUFA supports B cell metabolic fitness. (A) Flow cytometry of analysis of murine B cell proliferation and class switch to IgG1. Purified splenic B cells were activated with LPS/IL-4 in presence of vehicle (BSA), OA alone (100 μM), etomoxir (40 μM) and OA plus etomoxir at 72 h. Right, summary of IgG1^+^ B cell percentage normalized to vehicle treated sample. (B) Oxygen consumption rate (OCR) was measured with a Seahorse XFe96 analyzer using CpG/IL-4/IL-5 activated murine B cells in the presence of BSA, OA, PO, SA and PA for 48 h. Right, summary of basal respiration. (C) Measurement of OCR in human B cells activated in the presence of BSA, OA and PO for 72 h. Right, summary of basal respiration respiration. (D) Basal respiration was calculated in activated murine B cells with BSA and titrated doses of exogenous OA (1, 10, 20 and 50 μM). (E) Glycolytic capacity of activated murine B cells was measured in presence of BSA, OA, PO, SA and PA at 48 h using a Glycolysis Stress test. Right, summary of glycolytic capacity. (F) Extracellular acidification rate (ECAR) was measured in activated human B cells in the presence of BSA, OA and PO for 72 h. Right, summary of basal glycolysis rate. (G) Measurement of OCR and ECAR in the activated murine B cells in the presence of vehicle or SSI-4 for 48 h. (H) Staining of TMRM in activated murine B cell in the presence of vehicle, SSI-4, SSI-4 plus OA and OA alone for 72 h. Right, numbers indicate the MFI of TMRM. *p* values were calculated with one-way ANOVA. NS, not significant. Data were representative of 3 (A and H) or 2 (B, C, E and G) independent experiments. Error bars represent SEM.

### SCD-mediated MUFA supports mTORC1 activity and prevent excessive autophagy

SCD mediated FA metabolism has been linked to autophagy induction in fibroblasts and cancer cells, but there are contradictory findings^24–26^. It is unclear how FA metabolism may link to autophagy in lymphocytes. To examine how SCD inhibition affects subcellular organelles, we performed transmission electron microscopy on SSI-4 treated B cells, as well as SCD deficient B cells. We observed increased structures with double layer membrane encompassing various organelles or multi-lamellar membrane structures, which are signatures of autophagosomes^27^ (Fig. 4A and Fig. S4A). The conversion of the soluble form of LC3 (LC3-I) to the autophagic vesicle-associated form (LC3-II) is a hallmark of autophagy. Immunoblot analysis showed that SSI-4 treated B cells (Fig. 4B) and SCD deficient B cells (Fig. 4C) had substantial increase of LC3-II/LC3-I ratio compared to control cells. Treatment with exogenous OA completely rescued the increased LC3-II/LC3-I ratio (Fig. 4B, 4C). Consistent with the immunoblot data, B cells isolated from GFP-LC3 reporter mouse had higher LC3 expression after inhibition of SCD activity by SSI-4 *ex vivo* (Fig. 4D). It is well established that mTORC1 controls autophagy and cell metabolism^28^. To examine whether the mTOR pathway is involved in the inhibition of SCD activity induced autophagy, we analyzed the phosphorylation levels of S6K, and S6, which are the direct downstream substrates of mTORC1 pathway. Our results showed that the levels of p-S6K and p-S6 were decreased in SSI-4 treated B cells (Fig. 4B) or SCD1/2 deficient B cells (Fig. 4C), which were restored by exogenous OA (Fig. 4B, 4C). Reduced expression of activation-induced cytidine deaminase (AID) was also observed upon SSI-4 treatment, consistent with the positive effect of mTORC1 on AID induction^10^ (Fig. 4B). Furthermore, SCD inhibition also suppressed mTORC1 activity in B cells *in vivo*, when mice were fed with SSI-4 chow or control chow followed by immunization with hapten 4-hydroxy-3-nitrophenylacetyl conjugated with ovalbumin (NP-OVA) (Fig. 4E). Furthermore, we tested whether suppression of autophagy can rectify the B cell defects *in vitro*. Treatment with classic autophagy inhibitors, 3-methyladenine (3-MA) and wortmannin, both targeting class III PI3K, successfully rescued the B cell proliferation and class switch to IgG1 in the presence of SSI-4 (Fig. 4F). Lastly, increased SFA content is associated with increased endoplasmic reticulum (ER) stress^29^. Because SCD inhibition increases the SFA/MUFA ratio (Fig. 1F), it could also induce ER stress. Indeed, we found that the ER stress-related genes, including *Chop, Atf4*, and *Sqstm1*, were increased after SSI-4 treatment, while exogenous OA reduced these genes upregulation (Fig. S4B). Taken together, these data showed that SCD-mediated MUFA availability is required for suppressing ER stress and maintenance of mTORC1 activity, which prevents overactivation of autophagy.

**Figure 4.**
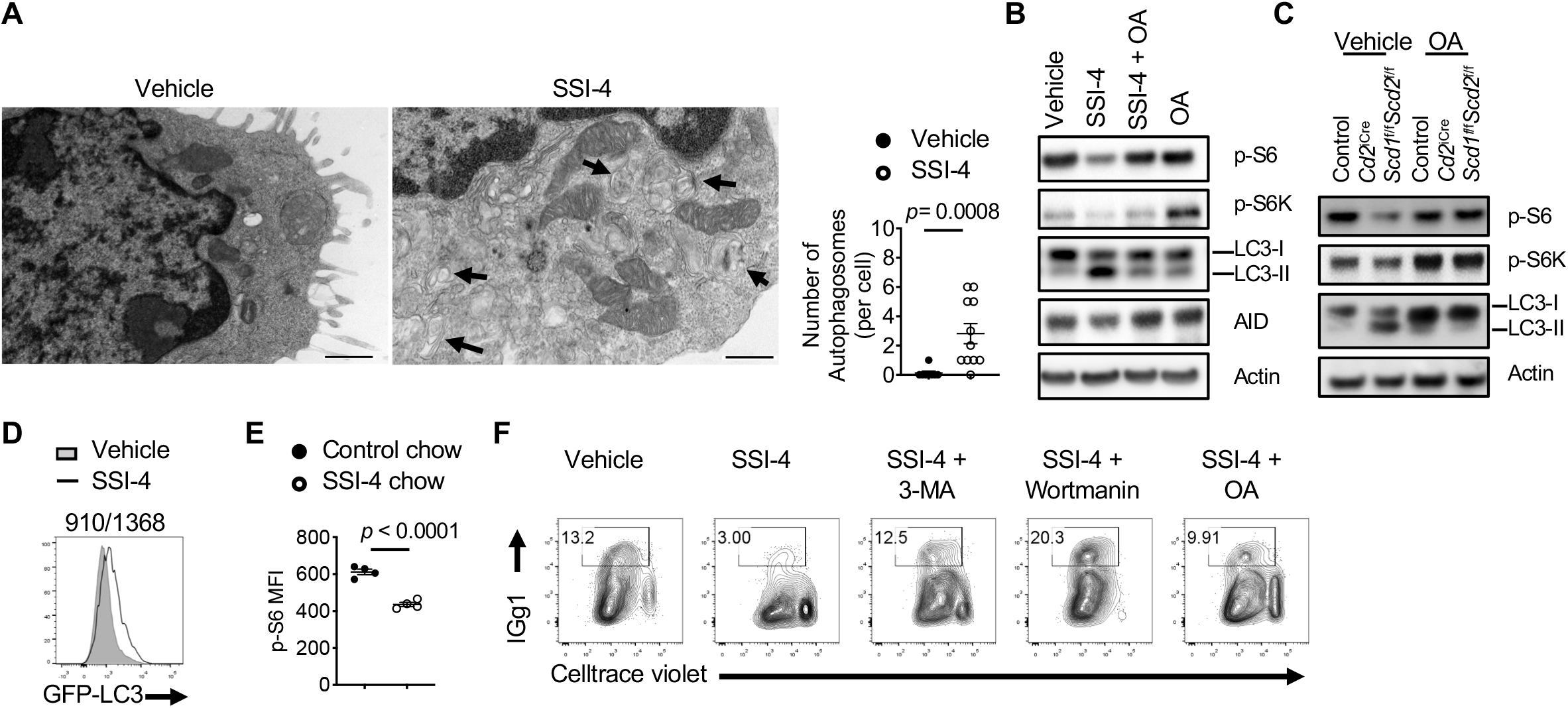
Inhibition of SCD activity induces the formation of autophagosomes and suppresses mTORC1 activity. (A) Transmission electron microscope image of activated mouse B cell treated with vehicle or SSI-4. Arrows indicate autophagosome structures. Right, numbers of autophagosomes found in vehicle (n = 8) or SSI-4 (n = 11) treated B cells. Scale bars: 0.5 μm. (B) Immunoblot analysis of LC3-I/LC3-II, p-S6, p-S6K, and AID in activated murine B cells in presence of vehicle, SSI-4, SSI-4 plus OA and OA alone. (C) Immunoblot analysis of LC3-I/LC3-II, p-S6, and p-S6K in activated B cells from *Cd2^iCre^Scd1*^+/+^*Scd2*^+/+^ and *Cd2^iCre^Scd1*^fl/fl^*Scd2*^fl/fl^ mice with or without exogenous OA. (D) Flow cytometry analysis of GFP-LC-3 expression in activated B cells isolated from GFP-LC-3 reporter mouse treated with vehicle, and SSI-4. Numbers indicate the MFI of GFP. (E) Mice were fed with control chow and SSI-4 chow (30 mg/kg), followed by NP-OVA immunization for one week. MFI of p-S6 in splenic B cells was measured by flow cytometry. (F) Cell proliferation and class switch recombination in activated murine B cells treated with SSI-4 in the presence of 3-methyadenine (3-MA) (1 mM) and wortmannin (1 μM), or OA. *p* was calculated with Student’s *t*-test. Data were representative of 2 (B and C) and 3 (F) independent experiments. Error bars represent SEM.

### Systemic SCD activity supports early B cell development and humoral immune response in vivo

Next, we investigated how SCD activity contributes to B cell development and function *in vivo*. Feeding mice with SSI-4 chow for 1 week markedly reduced serum OA and PO concentration, demonstrating the efficacy of SSI-4 *in vivo* (Fig. S5A). At this time, SSI-4 treatment resulted in significant reduction of the pool of immature B cells and B cell precursors (CD19^+^B220^int^) in bone marrow (BM) (Fig. 5A) and CD25^+^ pre-B cells (Fig. 5B). The composition of thymocytes were unaffected after SSI-4 treatment (Fig. S5B). In addition, the number of peripheral B cells and CD4^+^ T cells were not affected by the reduced OA content (Fig. 5C-5E), although the frequency of mature B cells (CD19^+^IgM^lo^IgD^+^) was slightly increased with SSI-4 chow treatment (Fig. S5C). Moreover, we observed a modest reduction of GC percentage and IgA expression in Peyer’s patches, where gut microbiota drives constitutive generation of GC and class switch to IgA^30^ (Fig. S5D, S5E), suggesting that SCD activity may be required for these processes. We next evaluated the impact of SCD inhibition on humoral immune response upon stimulation with foreign antigen, NP-OVA, *in vivo*. Notably, we observed that immunization increased OA concentration in sera (Fig. 5F), indicating that immune challenge is associated with increase of systemic MUFA content *in vivo*. SSI-4 treatment substantially reduced the OA level in immunized mice as expected (Fig. 5G). Importantly, SCD inhibition significantly suppressed splenic GC formation (Fig. 5H) and reduced antigen specific NP^+^ GC B cells (Fig. 5I). However, B220^int^CD138^+^ plasmablast generation was not affected by inhibition of SCD activity, suggesting that GC formation, but not plasmablast differentiation, is particularly sensitive to MUFA availability (Fig. 5J). Furthermore, the percentage of Tfh cells after immunization was comparable between these two groups, suggesting that SCD activity preferentially supported B cell activation (Fig. 5K). Consequently, the production of anti-NP specific total IgG and IgG1, but not IgM, antibodies were significantly reduced in response to systemic inhibition of SCD activity (Fig. 5L). Therefore, SCD activity supports early B cell development in the BM, and promotes GC formation and antibody production upon immunization.

**Figure 5.**
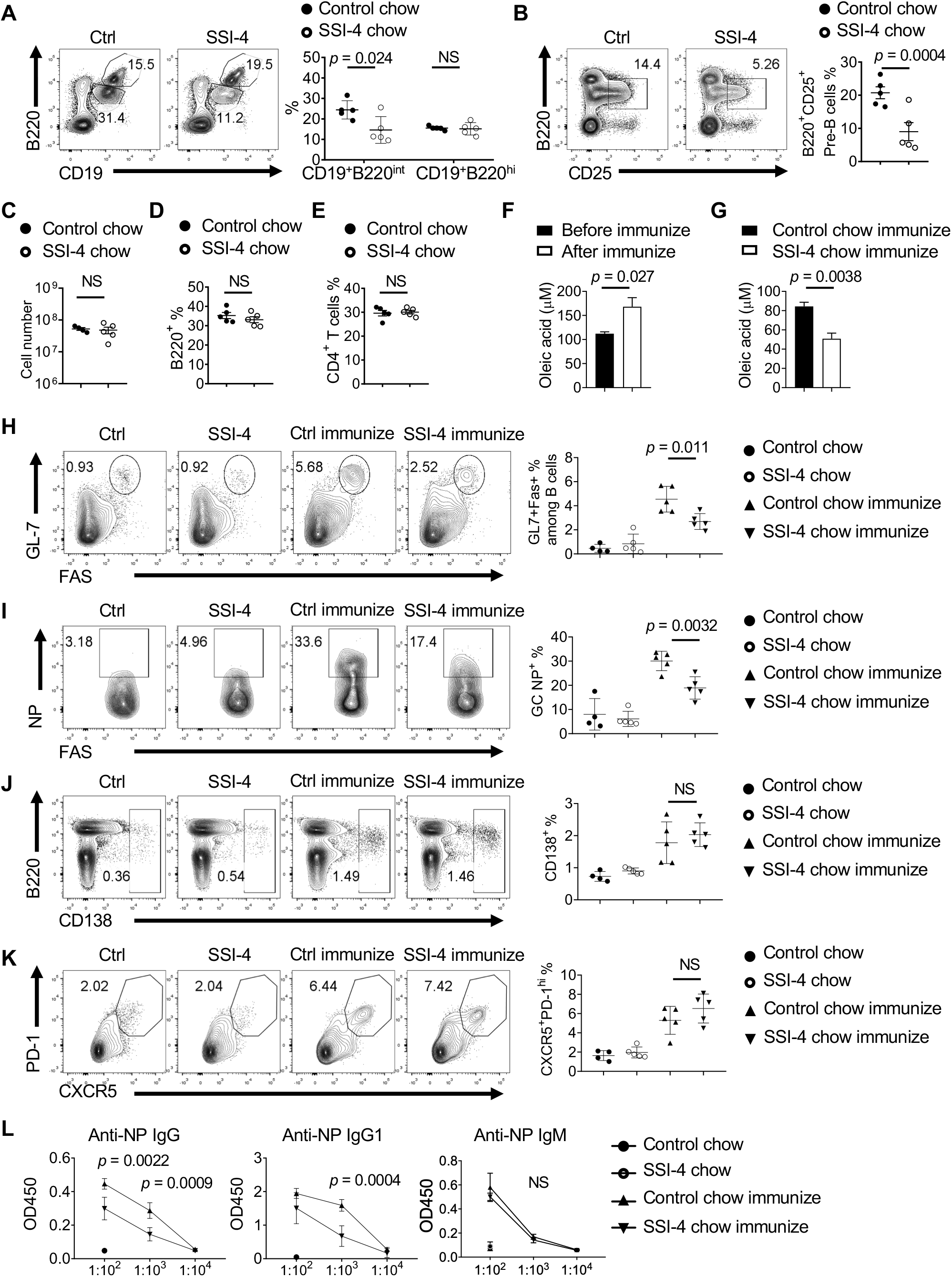
Systemic SCD activity is essential for humoral immune response *in vivo.* (A) Flow cytometry analysis of B cell precursors (B220^int^CD19^+^) and circulating mature B cells (B220^hi^CD19^+^) in bone marrow from mice fed with control chow or SSI-4 chow. Right, frequencies of B cell precursors and circulating mature B cells. (B) Flow cytometry of pre-B cell (B220^+^CD25^+^) in bone marrow from mice fed with control chow or SSI-4 chow. Right, frequencies of pre-B cells. (C-E) Mice were fed with control chow or SSI-4 chow for 2 weeks. The numbers of whole spleen (C), splenic B220^+^ B cells (D), and CD4^+^ T cells (E). (F) OA concentration in sera was detected by GC-MS before and after immunization. (G) OA concentration in sera was measured by GC-MS in mice immunized with NP-OVA and treated with control chow or SSI-4 chow. (H-J) Mice were divided into 4 groups. They were fed with control chow or SSI-4 chow, followed with or without NP-OVA immunization. Flow cytometry of germinal center (GC) (H), antigen specific NP^+^ expression among GC B cells (I), B220^int^CD138^+^ expression (J), and expression of PD-1 and CXCR5 among CD4^+^ T cell (K). Right, the frequencies of GC B cells (H), NP^+^ GC B cells (I), B220^int^CD138^+^ plasmablasts (J) and PD-1^+^CXCR5^+^ Tfh cells (K). (L) Measurements of anti-NP immunoglobulins in serial diluted serum from unimmunized and immunized mice fed with control chow or SSI-4 chow, presented as absorbance at 450 nM (A_450_) in ELISA. *p* values were calculated with Student’s *t*-test. NS, not significant. Data were representative of 2 (A-G) and 3 (H-L) independent experiments. Error bars represent SEM.

### SCD activity is required for humoral immunity against influenza infection

Humoral immunity plays a critical role against influenza infection^31^. How FA metabolism regulates anti-influenza immunity remains incompletely understood. Previous study showed that deficiency of FABP5 led to increased anti-influenza antibody production^32^. Deficiencies of FABP family molecules are associated with increased MUFA content in mice, suggesting that MUFA availability may contribute to anti-influenza humoral immunity^33,34^. Indeed, we observed increased serum OA and PO concentration, and OA/SA and PO/PA ratio after H1N1 influenza A infection, suggesting that influenza infection might trigger SCD activity (Fig. 6A). Feeding with SSI-4 chow led to substantially more severe weight loss in influenza A infected mice compared to control chow (Fig. 6B). The systemic inhibition of SCD activity by feeding with SSI-4 chow significantly reduced the frequency of GC B cells (Fig. 6C), and the expression of Bcl6, key transcription factor for GC B cell differentiation, in B cells (Fig. 6D). However, it did not affect CD138^+^ plasmablast formation (Fig. 6E), nor did it affect CXCR5^+^Bcl6^hi^ Tfh cell differentiation (Fig. 6F). Consequently, SCD inhibition significantly dampened the production of anti-influenza total IgG, IgG1 and IgG2c, but not IgM, in sera (Fig. 6G). Altogether, these data suggested that SCD activity is essential for antigen specific GC B cell formation and antibody production upon immunization and respiratory viral infection *in vivo*.

**Figure 6.**
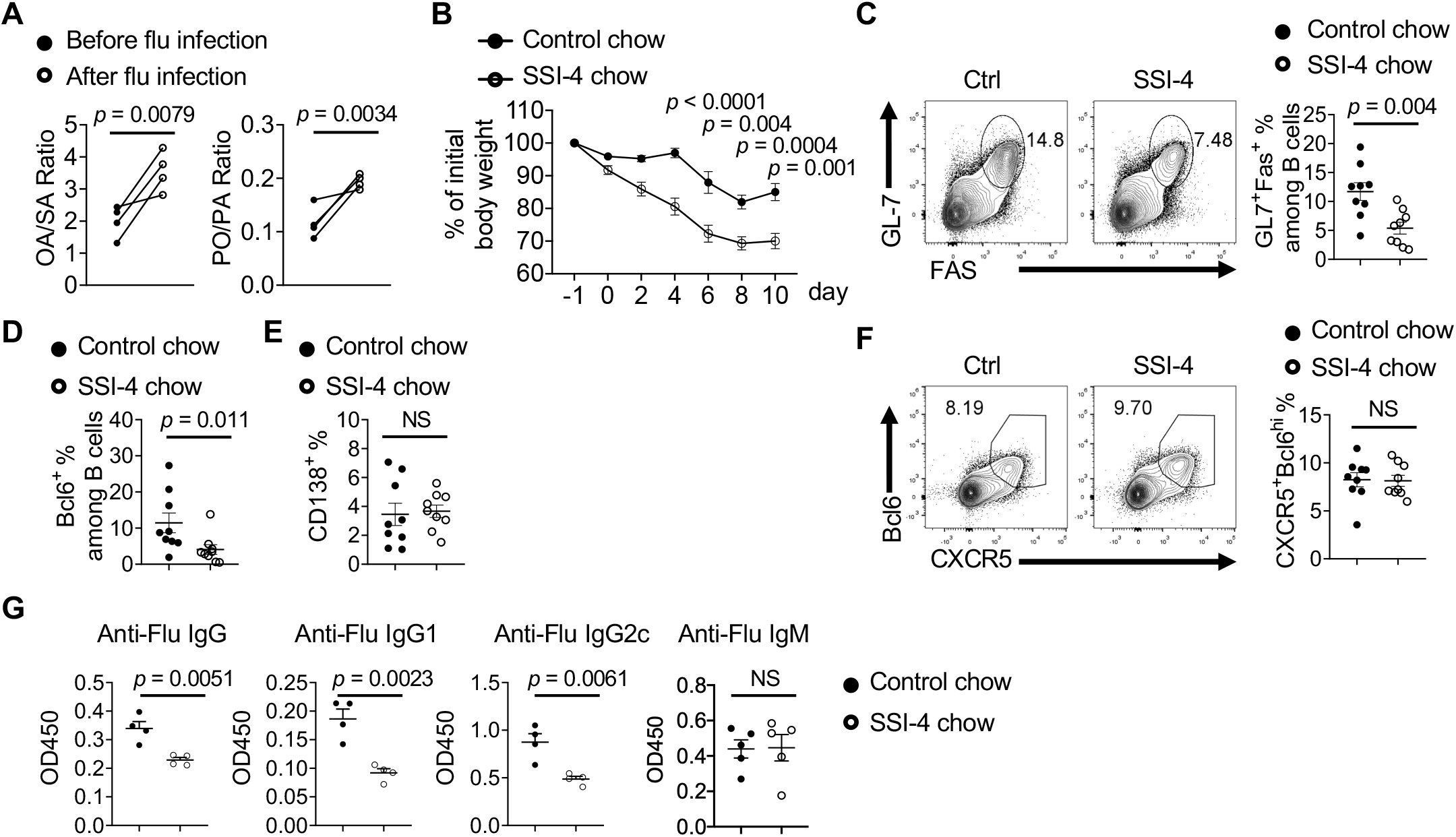
SCD activity is required for humoral immunity against influenza infection. (A) The ratio of OA/SA and PA/PO in sera before and after influenza infection. (B) Body weight change in mice fed with control chow or SSI-4 chow for one week following influenza infection. (C) Flow cytometry analysis of GC B cells in mediastinal lymph nodes at day 11 following infection. Right, the frequencies of GC B cells. (D) The frequencies of Bcl6^+^ expression among B cells in mediastinal lymph nodes. (E) The frequencies of B220^int^CD138^+^ plasmablasts. (F) Flow cytometry analysis of Tfh cells in mediastinal lymph nodes. Right, the frequencies of CXCR5^+^Bcl6^hi^ Tfh among CD4^+^ T cells in mediastinal lymph nodes. (G) Influenza virus specific antibody IgG, IgG1, IgG2c and IgM in sera were measured by ELISA. *p* was calculated with Student’s *t*-test and one-way ANOVA. Results were pooled from 2 (A-F) independent experiments. Error bars represent SEM.

### Intrinsic SCD activity is not required for B cell development and B cell response in vivo

Although our above data clearly demonstrated that SCD activity was critically required for B cell development and function, it was unclear whether B cell intrinsic or extrinsic SCD activity was responsible *in vivo*. Because our *in vitro* experiments demonstrated that exogenously supplemented OA was able to rescue the SCD deficient B cells, it is possible that B cell-extrinsic SCD activity could compensate the loss of SCD in B cells. To examine whether SCD activity in lymphocytes is required for humoral immunity, we examined B cell development in *Cd2^iCre^Scd1*^fl/fl^*Scd2*^fl/fl^ mice, in which *Scd1* and *Scd2* were deleted in all lymphocytes from common lymphocyte precursors. We did not observe significant alteration in early B cell development, suggesting that B cell intrinsic SCD activity was dispensable for early B cell development (Fig. 7A). Moreover, we examined *Cd19^cre^Scd1*^fl/fl^*Scd2*^fl/fl^ mice, in which *Scd1* and *Scd2* were deleted specifically and efficiently in B cells at a later stage^35^ (Fig. S6A). To examine whether B cell intrinsic SCD activity contributed to B cell activation *in vivo*, we immunized *Cd19^cre^Scd1*^fl/fl^*Scd2*^fl/fl^ mice with NP-OVA. We did not observe significant defect in GC formation (Fig. 7B). To eliminate any possible secondary effects elicited by chronic SCD deficiency during B cell development, we constructed chimera mice by transferring purified SCD deficient B cells from tamoxifen treated *Cre*^ER^*Scd1*^fl/fl^*Scd2*^fl/fl^ mice, together with CD4 T cells from OT-II transgenic mice and WT mice, into *Rag1^−/−^* mice, followed by immunization with NP-OVA^36^. Again, we did not observe any differences in terms of GC formation (Fig. 7C) and antibody production (Fig. 7D). Altogether, these data indicate that B cell intrinsic SCD activity is not essential for B cell development and function, or alternatively, B cell extrinsic SCD activity can compensate for the loss of SCD activity in B cells to support humoral immunity.

**Figure 7.**
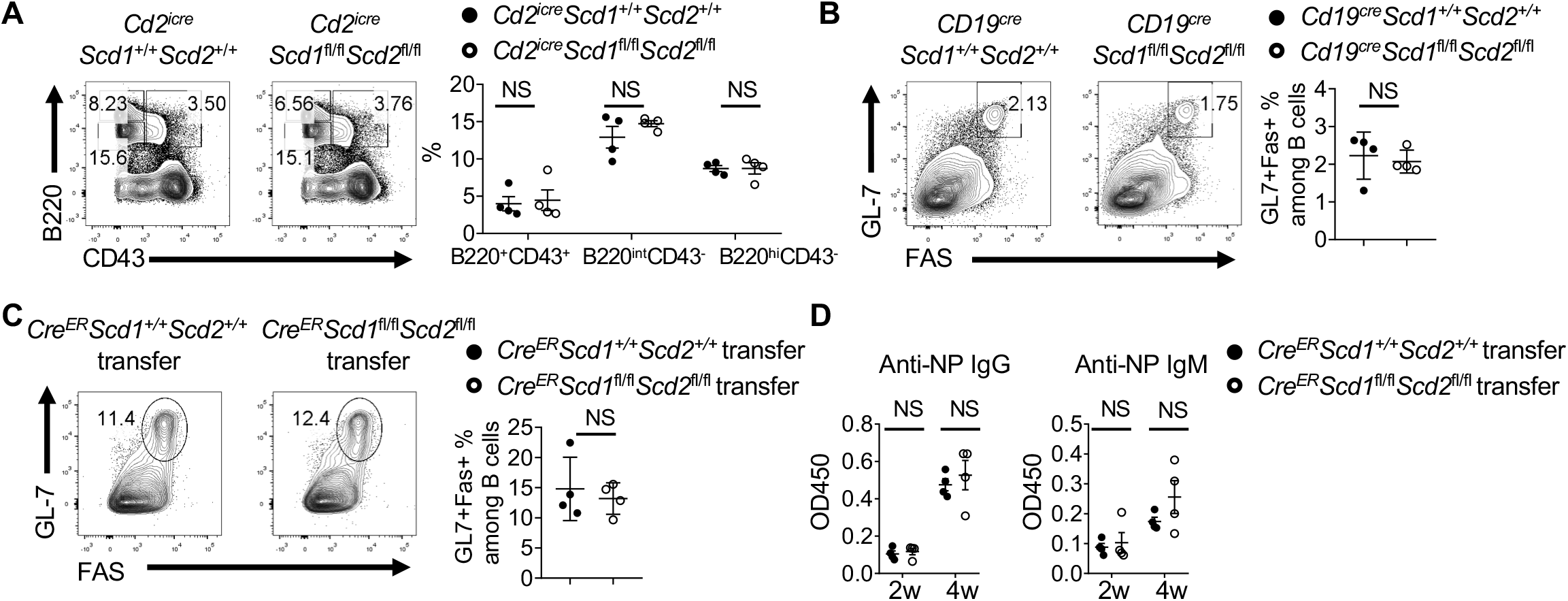
B cell intrinsic SCD activity is not required for B cell development and humoral response *in vivo.* (A) Flow cytometry analysis of B cell development in bone marrow from WT and *Cd2^iCre^Scd1*^fl/fl^*Scd2*^fl/fl^ mice. Right, the frequencies of B220^+^CD43^+^ pro-B cells, B220^int^CD43^−^ pre-B/immature B cells and B220^hi^CD43^−^ circulating mature B cells. (B) WT and *Cd19*^Cre^*Scd1*^fl/fl^ *Scd2*^fl/fl^ mice were immunized with NP-OVA. Flow cytometry analysis of GC B cells. Right, the frequencies of GC B cells. (C and D) B cells were purified from tamoxifen treated *Cre*^ER^*Scd1*^+/+^*Scd2*^+/+^ and *Cre*^ER^*Scd1*^fl/fl^*Scd2*^fl/fl^ mice. They were mixed with CD4 T cells purified from OT-II mice and WT C57BL/6 mice, and transferred into *Rag1^−/−^* mice. The recipient mice were immunized with NP-OVA. (C) Flow cytometry analysis of GC B cells in spleen. Right, the frequencies of GC B cells. (D) Measurements of anti-NP immunoglobulins in sera at 2 and 4 weeks after first immunization, presented as absorbance at 450 nM (A_450_) in ELISA. *p* was calculated with Student’s *t* test and one-way ANOVA. NS, not significant. Data were pooled from at least 3 (A) and represent 2 (C, D) independent experiments. Error bars represent SEM.

## DISCUSSION

The rapid progress in the immunometabolism field has revealed detailed mechanisms by which glucose and glutamine are metabolized during adaptive immune response, especially in T cell biology. However, our understanding of FA metabolism in lymphocytes remains highly limited. Historically, the impact of FAs on general immunity and inflammation has been a domain of nutrition science, and most of research in the past decades have been heavily focused on n-3 PUFA^37^. The impact of other FA species on adaptive immunity remains poorly defined. Our study offers a comprehensive study on the relationship between the availability of MUFA and humoral immunity. It reveals that murine and human B cells preferentially rely on OA to maintain their metabolic fitness and promote antibody production. SCD, which catalyzes the generation of endogenous OA, critically contributes to B cell development and activation. These observations are consistent with a recent study that GC B cells preferentially utilize FAO rather than glycolysis^14^. However, our results indicate that all FAs are not created equal for B cells. The quality of FA matters and proliferating B cell preferentially utilize MUFA than SFA. Thus, our data suggest that the SCD could constitutes a potential target for B cell mediated diseases.

SCD has been studied extensively for its function in various metabolic diseases, such as obesity, type 2 diabetes and hepatic steatosis. SCD1 deficiency protects mice from high fat diet or high carbohydrate induced obesity. The protection is partly attributed to hepatic SCD1 because deletion of SCD1 specifically in liver protected mice from high fat diet, but not high carbohydrate induced obesity^17,18^. It is proposed that SCD1 exerts its anti-adiposity function partly through suppressing FA oxidation and increasing FA biosynthesis genes^18,38^. SCD is also linked to cell autophagy, but there are conflicting evidence^24–26^. Our results demonstrated that SCD generated OA is required to sustain mTORC1 activity and mitochondrial metabolism in murine and human B cells. mTORC1 activity is critical for B cell isotype switch^10^ and mitochondrial metabolism supports B cell survival^5^. Thus, our results link availability of a MUFA to B cell antibody generation for the first time. In contrast to a recent study, we found no major defects in CD4^+^ T cell differentiation upon SCD inhibition, at least within our experimental schemes. The discrepancy could be due to the usage of different SCD inhibitors, different immune challenge models (primary intranasal infection using high-pathogenic PR8 strain vs intraperitoneal immunization using low-pathogenic X31 strain), and different time points of analysis^39^. Nevertheless, our results indicate that CD4^+^ T cells are less sensitive to SCD inhibition than B cells during respiratory influenza infection, and their metabolism may preferentially rely on different fuel sources compared to B cells, which warrant further investigation.

Notably, our data demonstrate that even though B cells have the capacity to synthesize MUFA, loss of B cell intrinsic SCD activity can be compensated by B cell extrinsic SCD activity, indicating that humoral immunity is supported by MUFA generated from non-lymphoid cells. While the source of the B cell-extrinsic SCD activity that sustains humoral immunity awaits future investigation, major metabolic organs expressing high level of SCD, such as liver and adipose tissues, are potential candidates^40^. Alternatively, stromal cells in lymphoid organs may also supply MUFA to support B cells at the local microenvironment. Recent studies in immunometabolism indicated that memory CD8^+^ T cells rely on intrinsically produced FAs to maintain their functions^41^. Our data showed that B cells may rely on extrinsic source of MUFA rather than synthesizing it themselves *in vivo*. Thus, B cells are nurtured by cell extrinsic FA and targeting non-immune cells may be required to modulating MUFA availability, and thus humoral immunity.

## MATERIALS and METHODS

### Isolation of human B cells

This study was conducted with approval from the institutional review boards of Mayo Clinic, Rochester. PBMCs were isolated from the medical waste following apheresis collection of platelets. Briefly, blood was diluted 1:3 using PBS. The diluted blood was then overlaid with Ficoll-Paque PLUS density gradient medium (GE Healthcare). The gradient was centrifuged at 400 g with no brake for 25 min at room temperature. The PBMC interphase layer was collected, washed with PBS with 0.1% BSA/2 mM of EDTA. Naive B cells (CD19^+^CD27^−^IgD^+^) were enriched using Human naive B cell negative selection kit (Stemcell Technologies).

### Mice

*Scd* floxed mice were from Dr. Makoto Miyazaki, University of Colorado School of Medicine. Mice were crossed with *Cre-ERT2*(REF^42^), *Cd2-iCre*^8^ *or Cd19-Cre*^43^ transgenic mice (Jackson Laboratory). C57BL/6 and *Rag1*^−/−^ mice were purchased from the Jackson Laboratory. Spleen cells from GFP-LC3 reporter mice were a generous gift from Dr. Douglas Green, St. Jude Children’s Research Hospital. For SCD activity in vivo inhibition experiment, animals were maintained on chow containing SCD inhibitor SSI-4 (30 mg/kg) or control chow, which were gifts from Dr. John A. Copland, III. The composition of other nutrients, vitamins, and minerals were equivalent between theses diets. After SSI-4 or control chow treatment for two weeks, mouse sera were collected, and the mice were sacrificed and the bone marrow, thymus, spleen and mesenteric lymph nodes were examined. One day prior to immunization, mice were fed with SSI-4 and control chow. Antigen for immunization was prepared by mixing NP-OVA (Biosearch Technologies), 10% KAL(SO_4_)_2_ dissolved in PBS at a ratio of 1:1, in the presence of LPS (*Escherichia coli* strain 055: B5; Sigma) at pH 7^44^. Mice were immunized intraperitoneally (100 μg NP-OVA and 10 μg LPS precipitated in alum) for analysis of GC B cell generation in spleen and NP-specific antibody response in serum. Transfer model were generated by transferring 5 × 10^6^ B cells isolated from *Cre*^ER^*Scd1*^fl/fl^*Scd2*^fl/fl^ or *Cre*^ER^*Scd1*^+/+^*Scd2*^+/+^ treated with tamoxifen for 4 consecutive days, mixing with 4 × 10^6^ CD4 T and 1 × 10^6^ OT-II transgenic T cells, into *Rag1*^−/−^ mice. Three weeks after first immunization, mice were boosted with NP-OVA. One week after second immunization, the mice were sacrificed, and the spleens were examined.

For influenza virus infection, influenza A/PR8 strain (200 pfu/mouse) were diluted in FBS-free DMEM media on ice and inoculated in anesthetized mice through intranasal route. Sera were collected before and two weeks after infection for fatty acid component measurement. The mice were fed with control chow or SSI-4 chow for 1 week following influenza infection before switching to regular diet. The mediastinal lymph nodes were analyzed for GC B cell formation and Tfh differentiation. Mice were bred and maintained under specific pathogen-free conditions in Department of Comparative Medicine of Mayo Clinic with IACUC approval.

### RNA-seq

RNA from isolated fresh B cells and B cells activated with LPS/IL-4 was extracted using a RNeasy kit (Qiagen) following the manufacturer’s instructions. After quality control, high quality total RNA was used to generate the RNA sequencing library. Paired-end RNA-seq reads were aligned to the mouse reference genome (mm10) using a spliced-aware reads mapper Tophat2 (v2.0.6)^45^. Pre- and post-alignment quality controls, gene level raw read count and normalized read count (i.e. FPKM) were performed using RSeQC package (v2.3.6) with NCBI mouse RefSeq gene model^46^. Differential gene expression analyses were conducted using edgeR (version 3.6.8) and the built-in “TMM” (trimmed mean of M-values) normalization method were used^47^. The criteria for selection of significant differentially expressed genes were: | log2 fold change | >= 1.0 and *p* value <=0.001.

### Immune cell Purification and Culture

Mouse B cells were isolated from pooled single cell suspensions of spleen and peripheral lymoh nodes using CD19 Microbeads (Miltenyi) or Mouse B cell Isolation Kit (Stemcell Technologies). B cells were stimulated with LPS (10 μg/mL; Sigma-Aldrich) and recombinant IL-4 cytokine (10 ng/mL; Tonbo Bioscience) with or without SCD inhibitor SSI-4, and proliferation was measured by dilution of CellTrace Violet proliferation dye (Thermo Fisher Scientific). B cells were also stimulated with anti-IgM (10 μg/mL; Jackson ImmunoResearch), anti-CD40 (10 μg/mL; BioXcell) and IL-4 (10 ng/mL; Tonbo Bioscience), or TLR ligand CpG (2.5 μM; IDT), recombinant IL-4 and IL-5 (10 ng/mL; Tonbo Bioscience) cytokines. To test the function of monounsaturated and saturated fatty acids, B cells were activated in presence of exogenous fatty acid-BSA conjugate. Palmitoleic acid (NU-CHEK PREP, INC), palmitic acid (NU-CHEK PREP, INC), stearic acid (NU-CHEK PREP, INC) were conjugated with fatty acid free BSA (Sigma-Aldrich) as previously described^48^. Human B cells were activated with CpG OND2006 (2.5 μM; IDT), recombinant human cytokine IL-10 (50 μg/mL; Peprotech), IL-15 (10 ng/mL; Peprotech), IL-4 (10 ng/mL; Biolegend), IL-2 (10 unit/mL) and anti-human CD40 (1 μg/mL; BioXcell). Human B cell proliferation was measured by ^3^[H]-thymidine incorporation (1 μCi/mL) (American Radiolabeled Chemicals).

### Non-esterified free fatty acids (NEFAs) and Total Fatty Acid Composition

Fatty acids and total fatty acid composition were measured against a standard curve on a triple quadrupole mass spectrometer coupled with an Ultra Pressure Liquid Chromatography system (LC/MS) as previously described. Briefly, the cell pellets were spiked with internal standard prior to extraction with tert-Butyl Methyl Ether (MTBE). Roughly 25% of the sample was dried down, hydrolyzed, re-extracted and brought up in running buffer for the analysis of total fatty acid composition. The remaining portion of the extract was dried down and brought up in running buffer prior to injecting on the LC/MS for the NEFA measurement. Data acquisition was performed under negative electrospray ionization condition.

### Mass spectrometer measurement of serum NEFA using liquid chromatography system

Fatty acids were measured against a standard curve on a Thermo Quantum Ultra triple quadrupole connected to a Waters Acquity Ultra high-pressure liquid chromatography system (LC/MS) as previously described. Briefly, 50 μl of serum was spiked with internal standard prior to extraction. The extracts were dried down and brought up in running buffer prior to injecting on the LC/MS. Data acquisition was performed under negative electrospray ionization condition^49^.

### NEFA isotopomer analysis

Briefly, 5 × 10^6^ activated B cells were washed with PBS and re-cultured in glucose-free medium RPMI medium containing 10% dialyzed FBS and uniformly labeled [^13^C]-glucose (2 g/L; Sigma-Aldrich) for 24 and 48 h. Cell pellets were lysed in 1 × PBS prior to lipid extraction. The extract was dried down and brought up in running buffer before underwent analysis on an Agilent 6550 iFunnel Q-TOF mass spectrometer/1290 Infinity liquid chromatographic system. A mixed standard containing 14 fatty acids was also run at the beginning and at the end of the sequence to generate retention time lock as well as unlabeled mass spectrum for each fatty acid. The mass spec was operating in negative electrospray ionization. Data was acquired in scan mode from 50-1700 m/z range. Data analysis was performed on Agilent Technologies software including Profinder, Mass Profiler Professional (MPP) and Vista Flux. Briefly, the features extracted from the data files were aligned using Profinder and converted to a compound exchange file (CEF) format to import in the MPP. The list was filtered for frequency and abundance to identify features that are present in all samples and with high abundance. A library was created using mass, molecular formula, and retention time of these features and used in conjunction with the Vista Flux software to determine the presence of isotopologue in individual feature. Fatty acids displaying a presence of isotopic pattern were annotated using retention time lock, accurate mass, and the METLIN database with an error of 5 ppm.

### GC-MS analysis of oleic acid and palmitoleic acid

Fatty acids were extracted from mouse serum samples using Lipid Extraction Kit (Biovison: K216). In brief, 25 μl of serum and 0.5 ml of Lipid Extraction Buffer containing 1 μg of D2-oleic acid (Cambridge isotope: DLM-689-0.1) and 1 μg of D14-palmitoleic acid (Cayman: 9000431) were mixed, vortexed for 1 min, and agitated for 15-20 min on an orbital shaker at room temperature. The samples were then centrifuged at 10,000 x g for 5 min and the supernatants containing lipids were dried under N_2_ gas. Dried metabolite samples were dissolved in 75 μL methoxyamine (20 mg/mL in pyridine) and incubated at 70 °C for 30 minutes. Samples were then further derivatized with 75 μL N-Methyl-N-(tert-butyldimethylsilyl) trifluoroacetamide (MTBSTFA) + 1% tertbutyldimetheylchlorosilane (TBDMCS) and incubated again at 70 °C for 30 minutes. Samples were analyzed using an Agilent 7890B GC coupled to a 5977A mass detector. 3 μL of derivatized sample were injected into an Agilent HP-5ms Ultra Inert column, and the GC oven temperature increased at 15 °C/min up to 215 °C, followed by 5 °C/min up to 260 °C, and finally at 25 °C/min up to 325 °C. The MS was operated in split-less mode with electron impact mode at 70 eV. Mass range of 50-700 was analyzed, recorded at 1,562 mass units/second. The following fatty acids were detected as TBDMS derivatives: oleic acid (m/z 339), D2-oleic acid (m/z 341), palmitoleic acid (m/z 311), and D14-palmitoleic acid (m/z 325). Data was analyzed using Agilent MassHunter Workstation Analysis and Agilent MSD ChemStation Data Analysis softwares. IsoPat^2^ software was used to adjust for natural abundance as previously performed^50^.

### Electron microscopy

Cells were fixed in Trumps fixative (pH 7.2) at 4°C overnight, spun down and the supernatant removed. They were re-suspended in 2% agarose which was allowed to cool and solidify. The cells in agarose were then post-fixed in 1% OsO4, dehydrated through a graded series of ethanols and embedded in Spurr resin. One hundred nm (or 0.1 mm) ultra-thin sections were mounted on 200-mesh copper grids, post-stained with lead citrate, and observed under a JEOL JEM-1400 transmission electron microscope at 80kV.

### Metabolic Assays

The bioenergetic activities of the OCR and ECAR were measured using a Seahorse XFe96 Extracellular Flux Analyzed following established protocols (Agilent). Briefly, B cells were seeded at 200,000 cells/well on Cell-Tak (Corning) coated XFe96 plate with fresh XF media (Seahorse XF RPMI medium containing 10 mM glucose, 2 mM L-glutamine, and 1 mM sodium pyruvate, PH 7.4; all reagents from Agilent). OCR was measured in the presence of Oligomycin (1.5 μM; Sigma-Aldrich), FCCP (1.5 μM; Sigma-Aldrich), and Rotenone (1 μM; Sigma-Aldrich)/ Antimycin A (1 μM; Sigma-Aldrich) in Mito Stress assay. For ECAR measurement, B cells were seeded in XFe96 plate with fresh Seahorse XF RPMI medium with 2 mM L-glutamine (PH 7.4), and treated with glucose (10 mM; Agilent), Oligomycin (1.5 μM), and 2-DG (50 mM; Sigma-Aldrich) orderly in Glycolysis Stress assay.

### Immunoblotting

For immunoblotting, cells were lysed in lysis buffer with protease and phosphatase inhibitors (Sigma-Aldrich). Protein concentration in samples were quantified by BCA assay (Thermo Fisher Scientific) before loading the samples for electrophoresis and membrane transfer. The transferred membrane was blocked with TBST (0.1% Tween 20) containing 5% BSA for 1 hr at room temperature. The membrane was incubated with primary antibodies overnight including anti-p-S6 (Ser235/Ser236, D57.2.2E; Cell Signaling), anti-p-p70 S6 kinase (Thr389, 108D2; Cell Signaling), anti-p-4E-BP1 (Thr37/46, 236B4; Cell Signaling), anti-LC3α/β (G-4; Santa Cruz); anti-p-ULK1 (Ser757, D7O6U; Cell Signaling), anti-AID (EK2 5G9; Cell Signaling), anti-β-actin (13E5; Sigma-Aldrich), and anti-SCD2 (H-12; Santa Cruz). Then, the membrane was washed and incubated with the corresponding secondary antibody for subsequent enhanced chemiluminescence (ECL; Thermo Fisher) exposure. The band intensity of all the immunoblot was analyzed by ImageJ software.

### Quantitative Real-time PCR

For mRNA analysis, total mRNA was isolated from mouse and human B cells by RNeasy Micro kit (Qiagen), reverse transcribed from mRNA to cDNA for subsequent real-time PCR analysis. *Scd1*, *Scd2, Fasn*, *Acaca*, *Chop*, *Atf4*, *Sqstm1* in mouse B cells and *Scd*, *Fasn*, *Acaca* expression in human B cells were measured by real-time PCR with a Bio-Rad Realtime PCR system. β-actin expression was used as control. The primers information was provided in the following table.

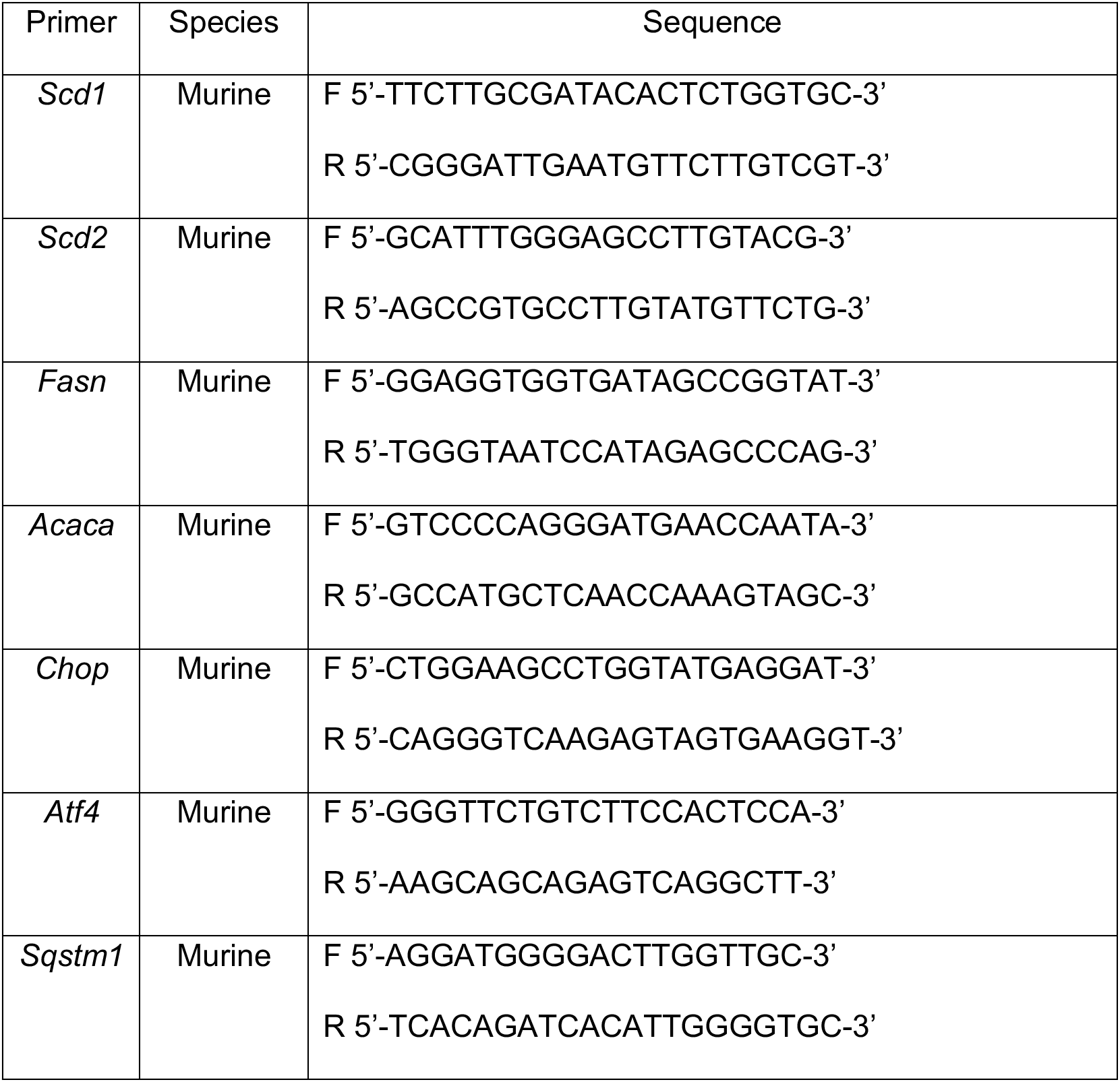
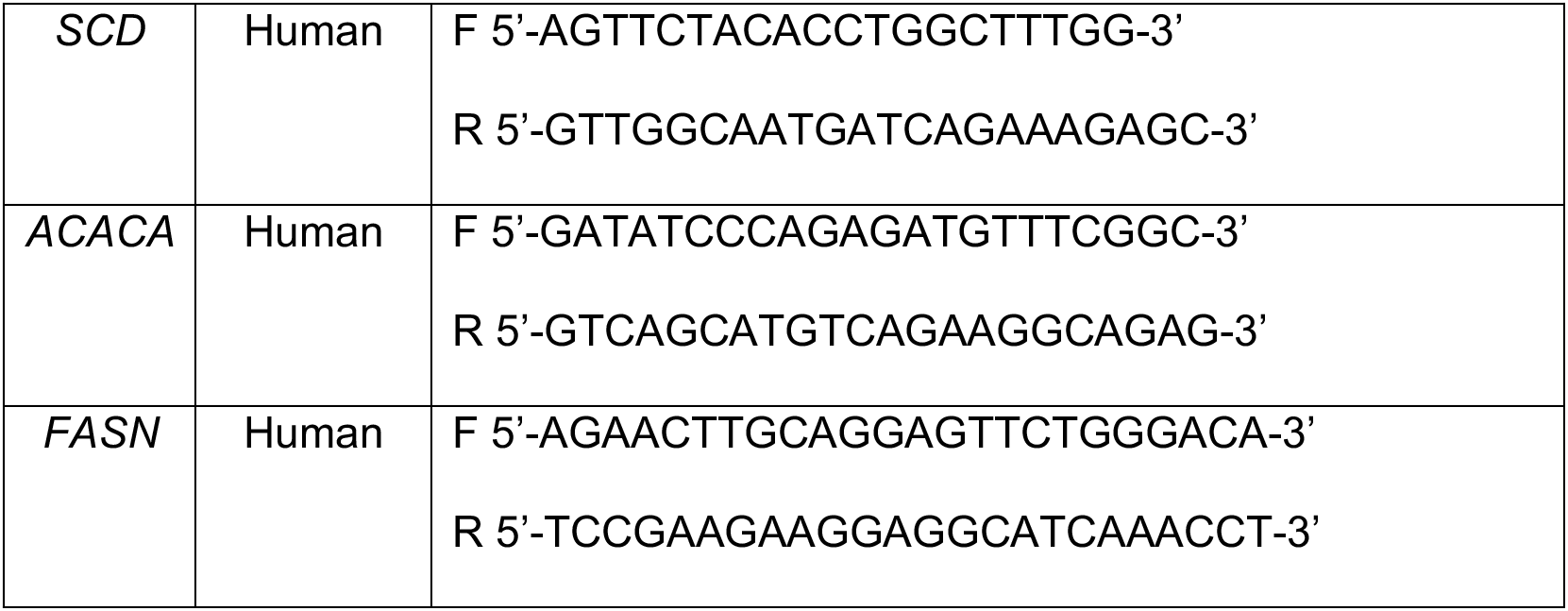

### Flow Cytometry

For analysis of surface markers, cells were stained in PBS containing 1% (w/v) bovine serum albumin on ice for 30 min, with anti-IgG1 (RMG1-1, Biolegend), anti-CD19 (ID3, Biolegend) anti-B220 (RA3-6B2, Biolegend), anti-CD4 (GK1.5, Biolegend), anti-CD8α (53-6.7, Biolegend), anti-CD25 (PC16, Biolegend), anti-GL7 (GL-7, Biolegend), anti-CD95 (Jo2, BD Biosciences), anti-CD138 (281-2, Biolegend), anti-IgD (11-26c.2a, Biolegend), anti-PD-1 (J43, ThermoFisher), anti-IgM (II/41, ThermoFisher). Antigen specific GC response was detected with tetramer NP-phycoerythrin conjugated with PE (Biosearch Technologies). CXCR5 was stained with biotinylated anti-CXCR5 (2G8) followed by staining with streptavidin-conjugated PE (both from BD Biosciences). Cell viability was assessed using the Fixable Dye Ghost 510 (Tonbo Bioscience) or Annexin V cell apoptosis kit with 7-AAD (ThermoFisher) following the manufacturer’s protocol. Phosflow staining for phospho-S6 (S235/236) was performed using Phosflow Fix/Perm kit (BD Biosciences). Mitochondrial potential was measured by staining with 20 nM TMRM (ThermoFisher) following manufacturer’s instructions. Flow cytometry was performed on the LSR II, LSR Fortessa (BD Biosciences) or Attune NxT (ThermoFisher) cytometers, and analysis was performed using FlowJo software (Tree Star).

## ELISA

For NP specific antibodies detection in sera, 96-well plates (2596; Costar) were coated with 2 μg/mL NP_23_-BSA in PBS overnight. Plates were washed twice (0.05% Tween 20 in PBS), blocked with 5% blocking protein (Bio-Rad) for 1 hr, and washed twice, and serially diluted serum samples were added for 1.5 hr at 37 °C. Plates were washed four times and horseradish peroxidase (HRP)-conjugated detection Abs for IgG (Bethyl Laboratories) and IgG1 (Bethyl Laboratories) were added for 1 hr at RT, washed four times, and tetramethylbenzidine (TMB) substrate was added. Reaction was stopped using 2N H_2_SO_4_ and read at 450 nm. Similarly, antibodies IgG, IgG1 and IgG2c specific to influenza A/PR8 strain in sera were measured with influenza virus coated plate.

### Statistical Analysis

Statistical analysis was performed using GraphPad Prism (version 8). P values were calculated with Student’s *t* test, or one-way ANOVA. *P* < 0.05 was considered significant.

## Acknowledgement

The authors thank Dr. Douglas Green at St. Jude Children’s Research Hospital for spleen cells from GFP-LC3 mice. We thank Dr. Michael Jensen for his expert advice and inputs on fatty acid metabolism. This work was partly supported by NIH R01 CA225680 (to T.H.), RO1 AG041756, RO1 AI112844 and RO1 AI147394 (to J.S.), Discovery Science Award from Center for Biomedical Discovery at Mayo Clinic (to H.Z.). Mayo Clinic Metabolomics Resource Core facility is supported by NIH U24DK100469.

---

**Supplementary Figure 1.**
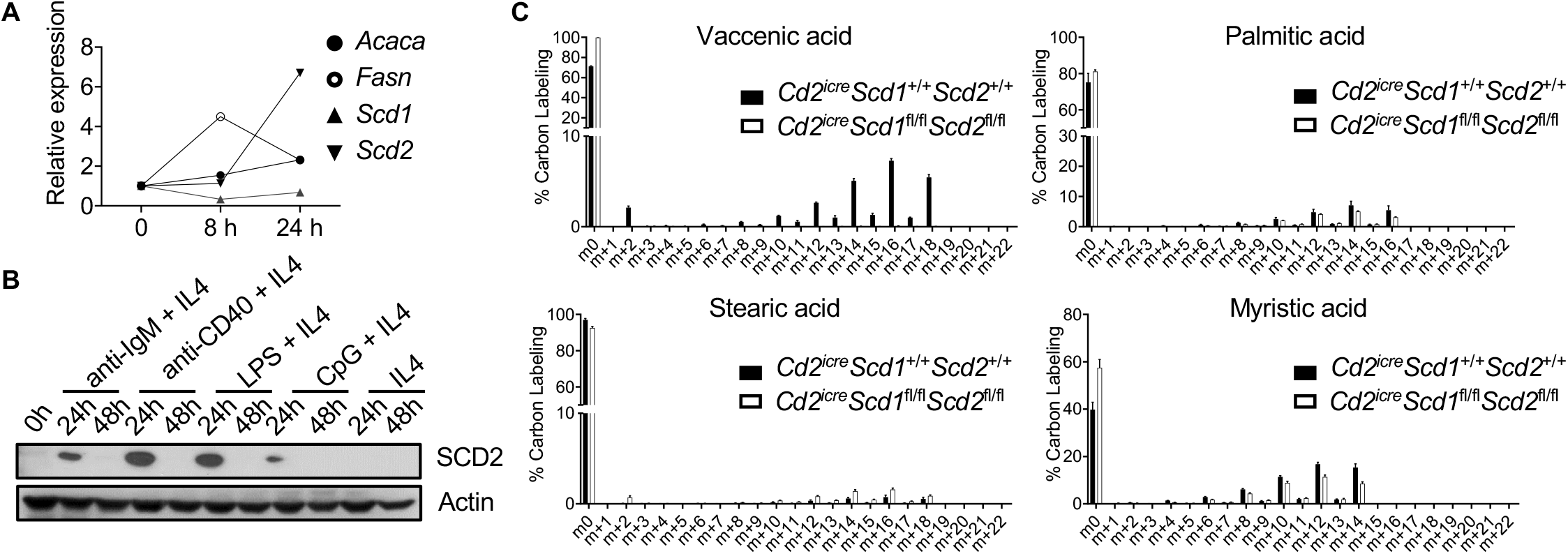
(A) Reverse transcription polymerase chain reaction (RT-PCR) analysis of *Acaca*, *Fasn*, *Scd1* and *Scd2* expression in murine B cells at 0, 8 and 24 h with LPS/IL-4 activation. (B) Immunoblot analysis of SCD2 in fresh isolated murine B cells, and B cells activated with anti-IgM (10 μg/mL)/IL-4, anti-CD40 (10 μg/mL)/IL-4, LPS (10 μg/mL)/IL-4, CpG (2.5 μM)/IL-4, or IL-4 alone at 24 and 48 h by immunoblot. (C) Incorporation of ^13^C labeled glucose into VA, PA, SA and MA were measured by LC-MS/MS in activated B cell isolated from WT or *Cd2^iCre^Scd1*^fl/fl^*Scd2*^fl/fl^ mice.

**Supplementary Figure 2.**
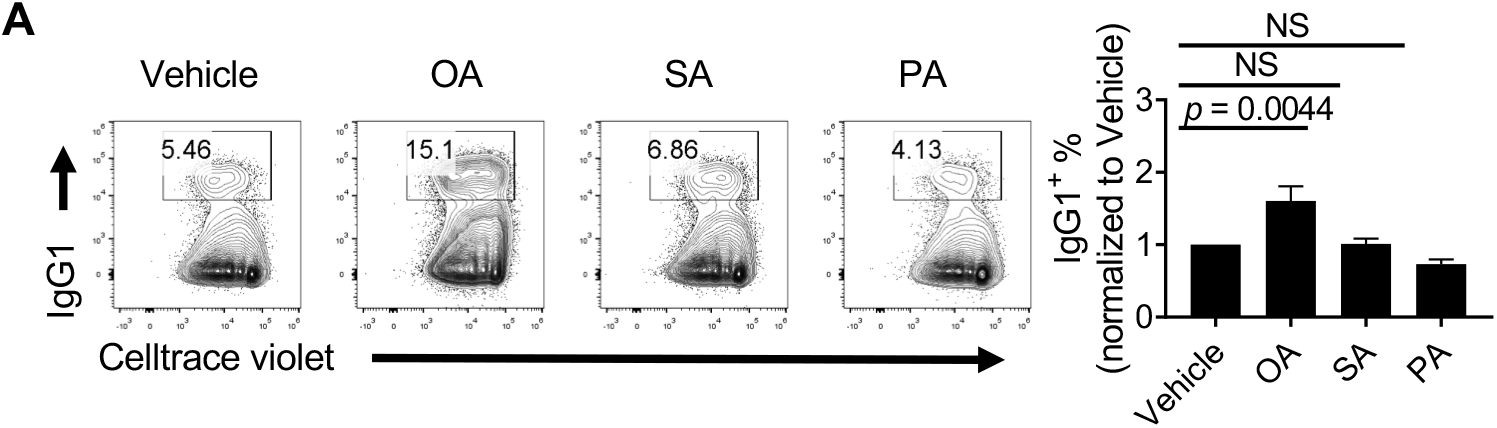
(A) Flow cytometry analysis of murine B cell proliferation and class switch in presence of OA, SA and PA. Right, the frequencies of IgG1^+^ B cell activated with LPS/IL-4 in presence of OA, PA and SA. The percentages were normalized to vehicle (BSA) group. *p* value was calculated with one-way ANOVA. NS, not significant. Results were pooled from 3 independent experiments. Error bars represent SEM.

**Supplementary Figure 3.**
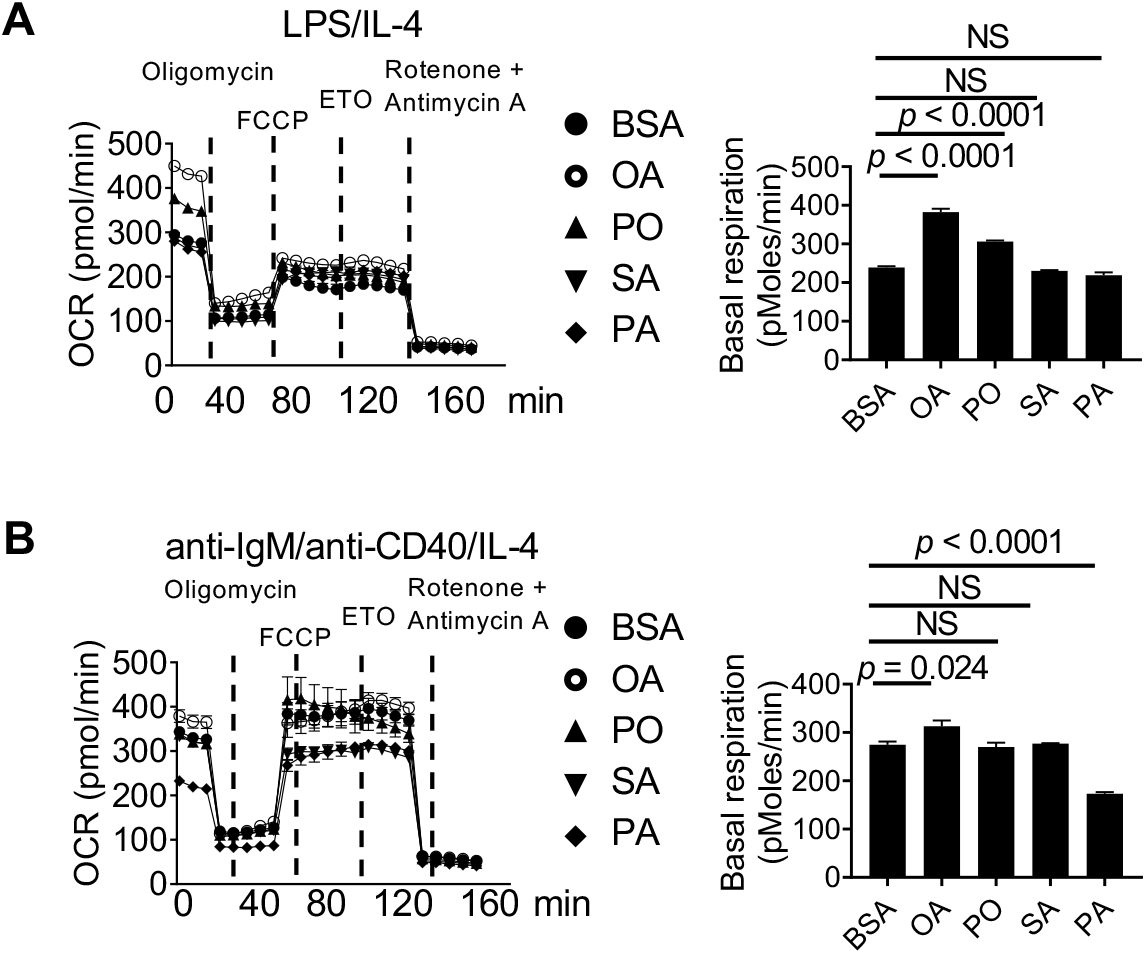
(A, B) Oxygen consumption rate (OCR) was measured with a Seahorse XFe96 analyzer using LPS/IL-4 (A) and anti-IgM/anti-CD40/IL-4 (B) activated murine B cells in the presence of BSA, OA, PO, SA and PA for 48 h. Basal respiration of either stimulation was summarized. *p* value was calculated with one-way ANOVA. NS, not significant. Data were representative of 2 (A and B) independent experiments. Error bars represent SEM.

**Supplementary Figure 4.**
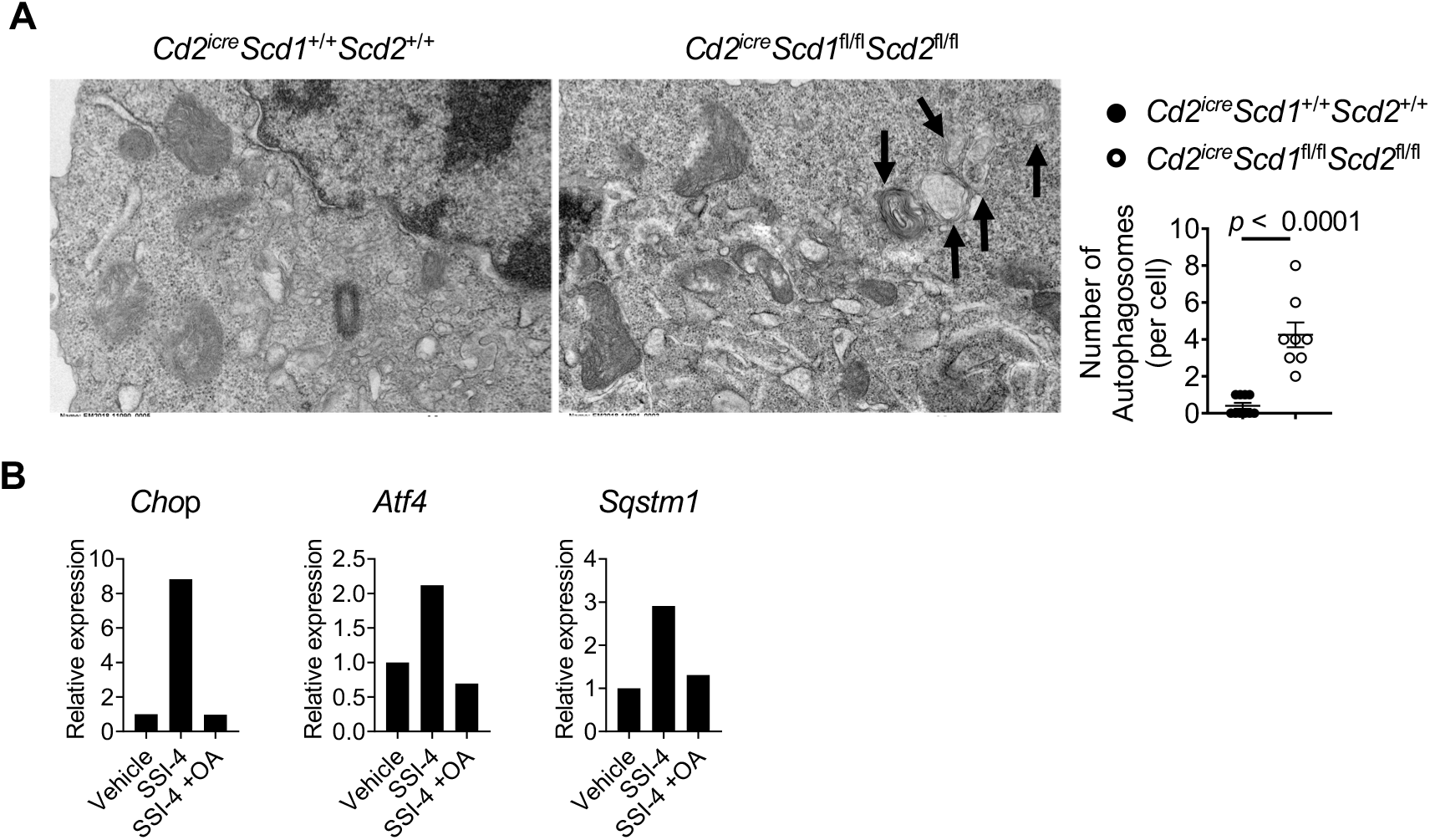
(A) B cells isolated from *Cd2^iCre^Scd1*^+/+^*Scd2*^+/+^ and *Cd2^iCre^Scd1*^fl/fl^*Scd2*^fl/fl^ mice were activated with LPS/IL-4 for 48 h. They were imaged on transmission electron microscope. Right, numbers of autophagosomes found in B cells from *Cd2^iCre^Scd1*^+/+^*Scd2*^+/+^ mouse (n = 10) and those from *Cd2^iCre^Scd1*^fl/fl^*Scd2*^fl/fl^ mouse (n =8). (B) Reverse transcription polymerase chain reaction (RT-PCR) analysis of *Chop*, *Atf4*, and *Sqstm1* in activated B cells in the presence of vehicle, SSI-4, or SSI-4 plus OA.

**Supplementary Figure 5.**
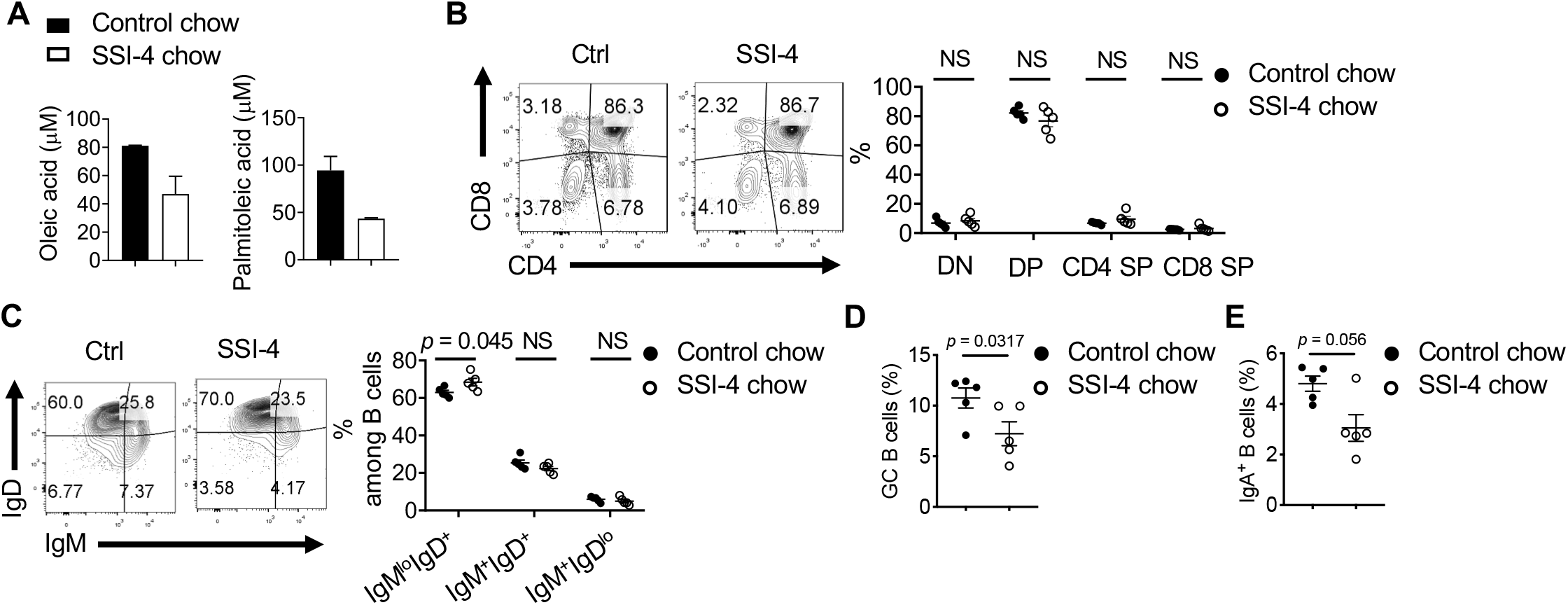
(A) OA and PO concentration in sera from mice fed with control chow or SSI-4 chow for 2 weeks. OA and PO were detected by GC-MS. (B) Flow cytometry analysis of thymic CD4^−^CD8^−^ (DN), CD4^+^CD8^+^ (DP), CD4^+^ single positive (SP) and CD8^+^ SP T cells from mice fed with control chow or SSI-4 chow. Right, the frequencies of DP, DN, CD4^+^ and CD8^+^ SP T cells. (C) Flow cytometry analysis of IgM^lo^IgD^+^, IgM^+^IgD^+^, and IgM^+^IgD^lo^ B cells among splenic CD19^+^ B cells. Right, the frequencies of each B cell subsets. (D-E) The frequencies of GC B cells (D) and IgA expression (E) among total B cells in Peyer’s patches from mice fed with control or SSI-4 chow. *p* was calculated with Student’s *t*-test. NS, not significant. Data were representative of 2 (A-E) independent experiments. Error bars represent SEM.

**Supplementary Figure 6.**
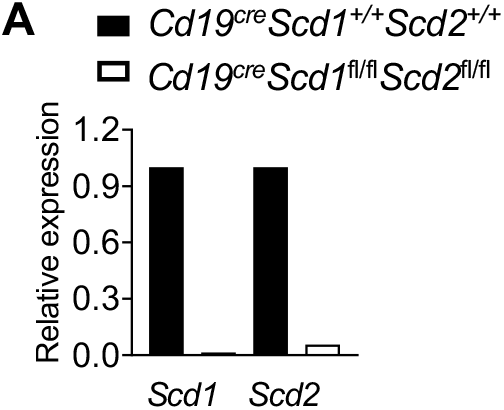
(A) RT-PCR analysis of *Scd1*, and *Scd2* expression in murine B cells isolated from WT and *Cd19*^Cre^*Scd1*^fl/fl^*Scd2*^fl/fl^ mice.

**Supplementary Table 1.** The concentrations of non-esterified free fatty acids (NEFA) were measured in female mice serum (8 weeks old, n = 8), including EPA, linolenic, DHA, myristic, palmitoleic, arachidonic, linoleic, palmitic, oleic, and stearic acids.

| Non-esterified free fatty acid (NEFA) | Concentration ($\mu\text{M}$, Mean ± SD) |
| --- | --- |
| EPA | 4.44 ± 0.56 |
| Linolenic acid | 44.6 ± 12.7 |
| DHA | 23 ± 3.5 |
| Myristic acid | 7.55 ± 2.17 |
| Palmitoleic acid | 31.7 ± 5.53 |
| Arachidonic acid | 14.8 ± 1.24 |
| Linoleic acid | 304 ± 74 |
| Palmitic acid | 239 ± 26.1 |
| Oleic acid | 184 ± 37.9 |
| Stearic acid | 53.8 ± 8.39 |

